# Loss of the centrosomal protein ALMS1 alters lipid metabolism and the regulation of extracellular matrix-related processes

**DOI:** 10.1101/2022.11.02.514847

**Authors:** Brais Bea-Mascato, Eduardo Gómez-Castañeda, Yara E. Sánchez-Corrales, Sergi Castellano, Diana Valverde

**Affiliations:** CINBIO, Universidad de Vigo, 36310 Vigo, Spain; Grupo de Investigación en Enfermedades Raras y Medicina Pediátrica, Instituto de Investigación Sanitaria Galicia Sur (IIS Galicia Sur), SERGAS-UVIGO, Vigo, Spain; Genetics and Genomic Medicine Department, Great Ormond Street Institute of Child Health, University College London, London, United Kingdom; Molecular and Cellular Immunology Section, Great Ormond Street Institute of Child Health, University College London, London, United Kingdom; UCL Genomics, Zayed Centre for Research into Rare Disease in Children, University College London, London, United Kingdom

**Author notes:** Correspondence: Diana Valverde., CINBIO Facultad de Biología, Universidad de Vigo, Campus As Lagoas-Marcosende s/n, 36310 Vigo, Spain.

**Keywords:** ALMS1, TGF-β, AKT, primary cilia, ciliopathy, ECM, lipid metabolism, Alström syndrome

## Abstract

**Background:** Alström syndrome (ALMS) is a rare autosomal recessive disease that is associated with mutations in *ALMS1* gene. The main clinical manifestations of ALMS are retinal dystrophy, obesity, type 2 diabetes mellitus, dilated cardiomyopathy and multi-organ fibrosis, characteristic in kidneys and liver. Depletion of the protein encoded by *ALMS1* has been associated with the alteration of different processes regulated *via* the primary cilium, such as the NOTCH or TGF-β signalling pathways. However, the cellular impact of these deregulated pathways in the absence of ALMS1 remains unknown.

**Methods:** In this study, we integrated RNA-seq and proteomic analysis to determine the gene expression profile of hTERT-BJ-5ta ALMS1 knockout fibroblasts after TGF-β stimulation. In addition, we studied alterations in cross-signalling between the TGF-β pathway and the AKT pathway in this cell line.

**Results:** We found that ALMS1 depletion affects the TGF-β pathway and its cross-signalling with other pathways such as PI3K/AKT, EGFR1 or p53. In addition, alterations associated with ALMS1 depletion clustered around the processes of extracellular matrix regulation and lipid metabolism in both the transcriptome and proteome. By studying the enriched pathways of common genes differentially expressed in the transcriptome and proteome, collagen fibril organisation, β-oxidation of fatty acids and eicosanoid metabolism emerged as key processes altered by the absence of ALMS1. Finally, an overactivation of the AKT pathway was determined in the absence of ALMS1 that could be explained by a decrease in *PTEN* gene expression.

**Conclusion:** ALMS1 deficiency disrupts cross-signalling between the TGF-β pathway and other dependent pathways in hTERT-BJ-5ta cells. Furthermore, altered cross-signalling impacts the regulation of extracellular matrix-related processes and fatty acid metabolism, and leads to over-activation of the AKT pathway.

## INTRODUCTION

Alström syndrome (ALMS, OMIM #203800) is a rare disease caused by mutations in the *ALMS1* gene. The subcellular localisation of the ALMS1 protein in the basal body of the primary cilium, composed of two centrioles, classified ALMS as a ciliopathy (Hearn et al., 2005; Knorz et al., 2010; Jagger et al., 2011; Kobayashi and Dynlacht, 2011). Ciliopathies are a group of diseases on which, the normal function and assembly of the primary cilia is affected. This ciliopathy is clinically characterised by the development of retinal dystrophy, type 2 diabetes mellitus (T2DM), obesity, dilated cardiomyopathy (DCM), hearing loss and multi-organ fibrosis affecting kidneys, liver and lungs (Collin et al., 2002; Zulato et al., 2011; Hearn, 2018).

Primary cilia are sensory organelles formed by microtubules protruding from the cell membrane. They are essential for a variety of physiological and developmental processes such as the control of cell cycle, migration and differentiation (Christensen et al., 2017; Pala et al., 2017; Anvarian et al., 2019). They also play a role in the regulation of various signalling pathways, such as WNT, Sonic Hedgehog (SHh), transforming growth factor β (TGF-β) and other G-protein-coupled receptor-regulated pathways (Ishikawa and Marshall, 2011; May-Simera et al., 2017, 2018; Pala et al., 2017; Anvarian et al., 2019).

The role of the ALMS1 protein is still unclear, with some (Graser et al., 2007; Li et al., 2007; Knorz et al., 2010) but not all (Collin et al., 2005, 2012; Hearn et al., 2005; Chen et al., 2017) studies suggesting a role in ciliogenesis. ALMS1 depletion, however, affects several signalling pathways regulated through primary cilia, such as the NOTCH and TGF-β pathways (Leitch et al., 2014; Álvarez-Satta et al., 2021; Bea-Mascato et al., 2022, 2023). In the NOTCH signalling pathway, ALMS1 depletion leads to the accumulation of receptors in late endosomes. However, it does not affect the recycling of these receptors in the cell line hTERT-RPE1 (Leitch et al., 2014). In the TGF-β pathway, inhibition of *ALMS1* expression lowers phosphorylation/activation of SMAD2 in hTERT-RPE1 (Álvarez-Satta et al., 2021), which could inhibit the canonical TGF-β pathway. Similar results have been described when abolishing other centrosomal proteins. For example, depletion of CEP128 decreases SMAD2 phosphorylation in zebrafish, hTERT-RPE1 and ciliated human foreskin fibroblasts (hFF) after TGF-β activation(Mönnich et al., 2018). However, these alterations were not observed in ALMS1 knockout models in HeLa or hTERT-BJ-5ta (Bea-Mascato et al., 2022). In these models the only alteration detected in the canonical pathway was a slight decrease in SMAD3 phosphorylation/activation in the hTERT-BJ-5ta model (Bea-Mascato et al., 2022). This suggests that alterations in the TGF-β pathway in the absence of ALMS1 may have a greater impact on other signalling pathways that cross-signal with this pathway such as PI3K/AKT or MAPKs (p38, c-JNKs and others) (Finnson et al., 2020).

Despite advances in recent years, little is known about the role of ALMS1 in primary cilia and its regulated signalling pathways. Here, we investigate at the RNA and protein level how the absence of ALMS1 expression in hTERT-BJ-5ta affects the TGF-β pathway and other cellular processes.

## MATERIALS AND METHODS

### Cell culture

hTERT-BJ-5ta human dermal fibroblasts and HeLa cell lines from American Type Culture Collection (ATCC) were used for this study. A 4:1 composite medium of Dulbecco’s minimum essential medium (DMEM, Gibco, Invitrogen, NY, USA) and Medium 199, Earle’s Salts (Gibco, Invitrogen, NY, USA), supplemented with 10% fetal bovine serum (FBS) (Gibco, Invitrogen, NY, USA) and 2% penicillin/streptomycin (P/S) (Gibco, Invitrogen, NY, USA) was used to maintain hTERT-BJ-5ta. DMEM medium supplemented with 10% FBS and 2% P/S was used for the HeLa cells. Both cell lines were cultured at 37°C with 5% CO_2_.

### RNA extraction

Initially, 3×10^5^ BJ-5ta cells were seeded in 6-well plates by triplicate in DMEM:199 10%FBS 2% P/S. After 24h a change of medium was made by adding DMEM:199 2% P/S without FBS and cells were incubated overnight for serum starvation. Next day, TGF-β pathway was stimulated by adding rhTGF-β1 ligand (2ng/mL; R&D Systems; 240-B) for 24 hours. Then, the medium was removed, and wells were washed twice with PBS. Cells were harvested in a tube of 1.5 mL after scraping them on PBS. The NYZ total RNA isolation kit (NYZtech, Lisboa, Portugal) was applied following the manufacturing protocol for RNA extraction. Finally, RNA was eluted, and sample concentrations were measured with nanodrop (Thermo Fisher, Waltham, USA).

### RNA-seq and library construction

RNA quality control was performed using Bioanalyzer (Agilent, Santa Clara, USA), finding a RIN ≥ 7. Then, RNA enrichment was carried out with the Dynabeads mRNA direct Micro kit (Life Technologies, Carlsbad, USA) following manufacturer’s protocol. Sequencing libraries were constructed from cDNA using the Ion Total RNA-seq Kit v2 with ERCC RNA Spike-in Control mix (Thermo Fisher, Waltham, USA). 25μL of each library was sequenced using a PI V2 chip in an Ion Proton Sequencer (Thermo Fisher, Waltham, USA).

After sequencing, fastq files were analysed in the Finisterrae II computer cluster of the “Centro de Supercomputación de Galicia” (CESGA). FastQC (http://www.bioinformatics.babraham.ac.uk/projects/fastqc) and MultiQC (Ewels et al., 2016) were used to determine the quality of the samples **(Figure S1 A)**. High-quality reads were aligned with STAR software (Dobin et al., 2013) using the primary assembly of Homo sapiens genome GRCh38.p13 (Gencode v32). We generated a count matrix using the HTSeq software (Anders et al., 2015). Downstream analysis was performed with the following R (version 4.0.5) packages: DESeq2 (Love et al., 2014) to detect differentially expressed genes (DEG; **Figure S1 B-D; Table S1**); and EnrichR (Kuleshov et al., 2016) for enrichment analysis of GO terms and Pathways, collected from different databases (Bioplanet, KEGG, MSigDB and wikipathway) **(Table S3)**. P-values were calculated using Fisher’s exact test and corrected using Benjamini-Hochberg (FDR). Finally, ggplot and pheatmap were used for the visualisation of the results (Wickham, 2016). To validate our homemade DESeq2 pipeline, SARTools (Varet et al., 2016) was applied and the same number of DEG was obtained.

The sequences generated in this work have been deposited in NCBI’s Gene Expression Omnibus (Barrett et al., 2013) and are accessible through GEO Series accession number GSE209844.

### Protein extraction

BJ-5ta cells were seeded at 3×10^5^ in 6-well plates by triplicate in DMEM:199 10%FBS 2%P/S. After 24 hours, FBS was removed from the medium and the cells were incubated overnight for serum starvation. The next day, cells were stimulated with rhTGF-β (2ng/mL) for 24 hours. After that, DMEM:199 was removed, and wells were washed twice with PBS. Cells were scrapped in PBS. Then, they were pellet at 10000 rpm for 5 min in a Sigma^®^ 1–14 K at 4°C and 100μL of RIPA buffer was added to each sample for cell lysis. Samples were incubated on ice for 10 min in constant agitation. Finally, cell debris was pelleted by centrifugation for 30 min at 12000 rpm and 4 °C. Samples were aliquoted and stored at −80 °C until the next step of the protocol. Protein quantification was performed by Bradford microplate assay using Bio-Rad protein assay (Bio-rad, Hercules, USA).

### Label-free proteomic profiling

Label-free proteomic profiling was performed following the same protocol described in (Bea-Mascato et al., 2022) with the corresponding quality controls **(Figure S2; Table S2)**. Ggplot and pheatmap were used for the results visualisation. Enrichment analysis was performed using the same protocol as described for the transcriptome **(Table S4)**. The mass spectrometry proteomics data of this study have been deposited to the ProteomeXchange Consortium via the PRIDE (Perez-Riverol et al., 2019) partner repository with the dataset identifier PXD035708.

### Networks generation

For the generation of the different networks, we used the programme Cytoscape (3.9.0) with Java (11.0.6). For the generation of the enriched route networks, the pipeline described by Reimand et al.(Reimand et al., 2019) was used. Similarity coefficients between nodes (minimum threshold of 0.375) were calculated using the arithmetic mean between the overlap coefficient and the Jaccard index of the different significantly enriched gene sets in the enrichment analysis (FDR < 0.05). After this procedure, the clusters were annotated using the plugins ClusterMaker2, wordCloud and autoanotation. The MCL algorithm was used via the ClusterMaker2 plugin to determine the clusters and then wordCloud and autoannotation were used to annotate the clusters. The settings were maintained to generate the networks of the different datasets.

### p-AKT/AKT activation assays

Initially, about 3×10^6^ BJ-5ta control and ALMS1 knockout (KO) cells were plated in 100mm dishes (Corning, NY, USA) in DMEM:199 2P/S% without FBS for serum starvation. After 24h of incubation, the rhTGF-β1 ligand was added at a final concentration of 2ng/mL for 0-, 10-, 30-, and 90-min. Dishes were then washed 3 times with PBS and harvested in 1mL of PBS, using a scraper. After that, cells were pelleted by centrifugation in a Sigma^®^ 1–14 K at 4°C, 7,400×g for 10 min and lysed on ice for 10 min using 200 μL of RIPA buffer, containing 1 mM sodium orthovanadate as phosphatase inhibitor (Sigma-Aldrich, Missouri, USA) and 0.1% (v/v) protease inhibitor cocktail (Merck, Darmstadt, Germany). To remove the pellet with cell debris not lysed with RIPA buffer, samples were centrifuged at 4°C, 14,500×g for 30 min, and the supernatant was collected in 1.5mL low binding tubes (Thermo Fisher, Waltham, USA). For protein quantification, the Bio-Rad protein assay reagent was used in a microplate assay. Finally, samples were stored at −80°C until analysis.

### SDS-PAGE, Western blot and quantification

Samples were prepared mixing 20μg of protein from each sample with 6.25μL of Laemmli Buffer 4X (Biorad, Hercules, USA), 1.25μL β-mercaptoethanol (BME) and H2Od up to a final volume of 25μL. After that, samples were boiled at 97°C for 5 min and run on a 12% Mini-PROTEAN^®^ TGX™ Precast Gel (Biorad, Hercules, USA) for 1 hour at 150V. Proteins were then transferred to a 0.2 μm polyvinylidene fluoride (PVDF) membrane using the Trans-Blot^®^ Turbo™ Transfer System (Biorad, Hercules, USA) following the mixed molecular weight (MW) protocol. Membranes were blocked with in-house TBS-T buffer [TBS/0.1% (v/v) Tween-20] with 5% (w/v) milk for 90min at room temperature (RT). Incubation with the primary antibody was carried out in blocking solution (TBS-T 5% milk) overnight at 4°C. For this assay, the following antibodies were used: anti-AKT (1:5000, Abcam, 179463) and anti-p-AKT (1:5000, Abcam, 222489).

The next day, the membranes were washed 3 times with PBS. They were then incubated with the secondary antibody in a blocking solution for 1 hour at RT. Goat anti-rabbit IgG H&L (HRP) (Abcam, 205718) was used as a secondary antibody in this assay at 1:5000 for p-AKT and 1:10000 for AKT.

Finally, Clarity western ECL substrate (Biorad, Hercules, USA) was used to develop the membranes. The photos were taken by exposure on the ChemiDoc system (Biorad, Hercules, USA) after 5 minutes of incubation with the substrate. Quantification of bands was carried on Image Lab software (Biorad, Hercules, USA) using the method of total protein.

### Fluorescent preparations

2×10^4^ HeLa cells (WT and *ALMS1* KO) were seeded in μ-Slide 8-well chambers (IBIDI, Germany). After 24 hours cells were transfected with pcDNA3 GFP-PTEN using Lipofectamine 3000 (Thermo Fisher, Waltham, USA) following the manufacturer’s protocol. Cells were incubated overnight at 37°C with 5% CO2. The following day, the medium was changed by adding DMEM 2% P/S and 2%FBS. In addition, rhTGF-β was added to the corresponding wells at a final concentration of 2ng/mL and the plates were incubated again overnight under the same conditions. Finally, on the fourth day, the preparations were marked, fixed and mounted, following the protocol detailed below.

200μL of DMEM medium per well containing CellMask (Thermo Fisher, Waltham, USA) at a 1:1000 dilution were added and incubated for 10min at 37°C in the dark. After two washes with PBS, the cells were fixed with 150 μL of 4% paraformaldehyde (PFA) in a 1:10 dilution of DMEM medium and incubated for 10min. The PFA diluted in DMEM was removed with 2 washes with PBS and 200 μL of PBS with DAPI was added at a final concentration of 1μg/mL, followed by a 20min incubation in the dark. Finally, the wells were washed 3 times with PBS to remove the DAPI solution and the samples were mounted with 3 drops of Prolong. After incubation for 1 hour at 4°C, the samples were analysed on the Nikon NIE microscope. (Nikon, Tokyo, Japan).

This experiment was performed in duplicate on 2 alternate days adding 2 wells in each experiment for each treatment and cell line (WT and KO).

GFP-PTEN was a gift from Alonzo Ross (Addgene plasmid # 13039; http://n2t.net/addgene:13039;RRID:Addgene_13039)

### Image acquisition, processing and analysis

The images were taken with the Nikon NIE direct microscope at 20X, using the RGB channels for the acquisition of DAPI (Blue), GFP-PTEN with FITC (Green) and CellMask with TRITC (Red). After that, all the images were processed with Fiji (Java 8 version) (Schindelin et al., 2012). The background was subtracted from all images, the resolution was set to 4908×3264 (0.25microns/pixel) and the RGB channels were grouped with the “Merge Channels” option to create the compositions, maintaining the independence of the channels. Cell segmentation was then performed using the biodock platform (https://www.biodock.ai/). Cells were filtered by size to remove any debris that could have been detected as cells by the algorithm. Finally, the signal from the 550 cells with the highest GFP intensity was normalised against the mean of TGF-β stimulated control to obtain the normalised fluorescence values.

### Statistical analysis

The proteomic study involved several statistical analyses and data transformations. The data were first normalized using a variance stabilisation procedure (VSN). Data imputation for missing values was carried out using the Minprob approach with a q-value of less than 0.01. Finally, differential expression analysis was performed using the linear protein models of the limma package (Ritchie et al., 2015) included in the DEP R package (Zhang et al., 2018).

Transcriptomic normalisation and analysis were performed using DESeq2 (Love et al., 2014). An FDR value < 0.05 was used to determine differentially expressed genes.

To analyse statistical differences in the Western blot data, a t-student with a Benjamini, Krieger, & Yekutieli two-stage correlation was used. Finally, the fluorescence imaging results were analysed with a 2-way ANOVA.

## RESULTS

### Inhibition of *ALMS1* gene expression causes several alterations of the transcriptome and proteome

In this study, we generated a multi-omics dataset (total RNA-seq and LFQ-proteomic analysis) on an hTERT fibroblast cell line (WT/CT and *ALMS1* KO). This cell line was previously stimulated with TGF-β1 ligand to identify alterations related with TGF-β pathway **(Figure 1A)**.

**Figure 1.**
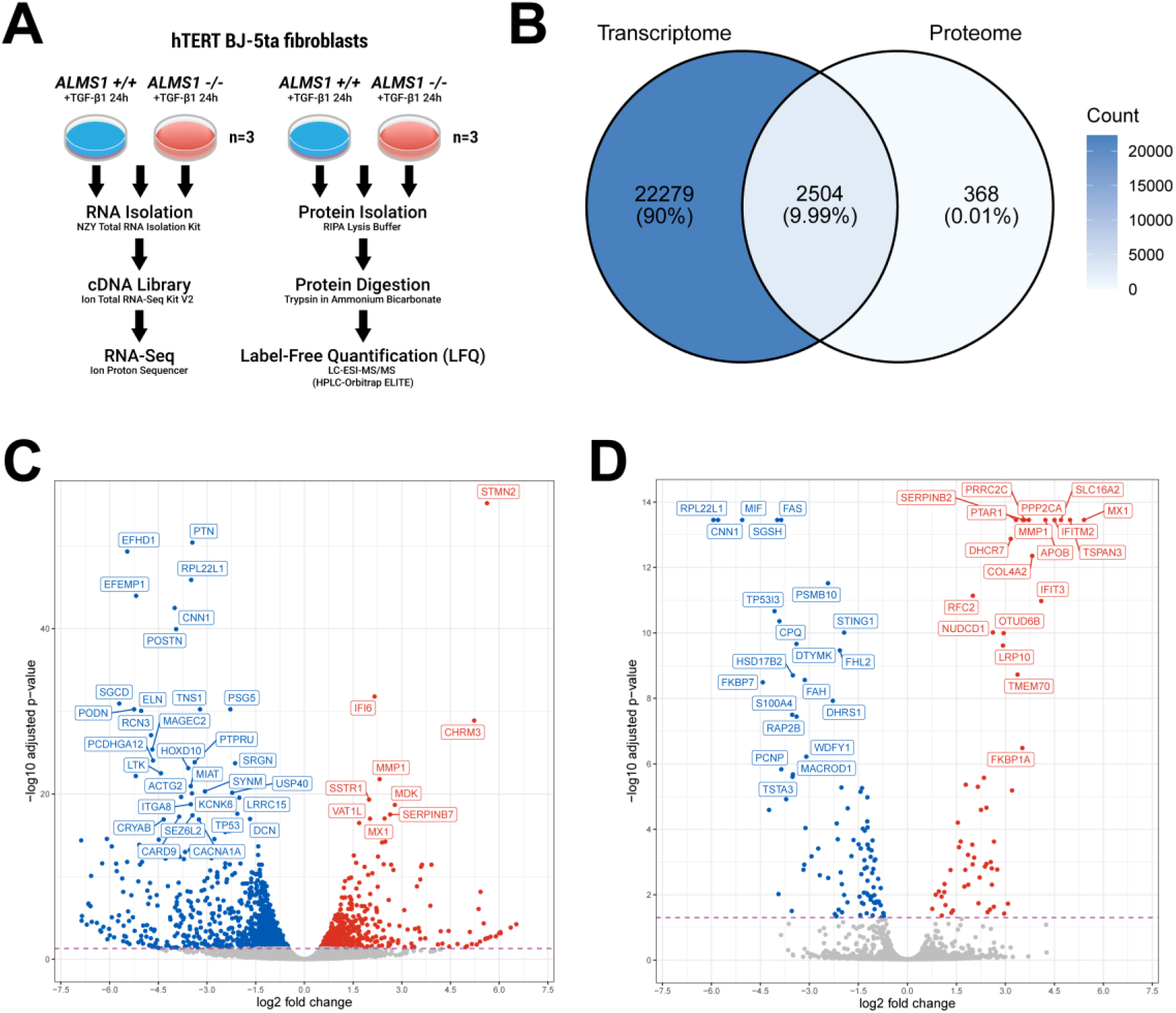
Differential expression analysis in the BJ-5ta ALMS1 knockout cell line by RNA-seq and LFQ-proteomics. **(A)** Illustration of the experimental design and procedure followed in this study**. (B)** Total overlap between genes identified in the transcriptome and the proteome. **(C)** Volcano-plot of differentially expressed genes (FDR < 0.05) in RNA-seq, highlighting the 40 most significant genes. **(D)** Volcano-plot of differentially expressed proteins (FDR < 0.05) in the proteome, highlighting the 40 most significant protein-coding genes.

We identified 24,784 genes (transcripts with distinct gene symbols) in the transcriptome **(Figure 1B, Table S1)**. On the other hand, we identified 2,872 proteins in the proteome, of which 1,884 were quantified in the 6 study samples (3WT and 3KO) **(Figure 1B, Figure S2A, B, Table S2)**. Of the proteins identified, 2,504 could be unambiguously associated with a gene. These protein-coding genes (PCG) identified in the proteome covered 10% of the total number of genes identified in the transcriptome **(Figure 1B).**

Our expression analysis (*ALMS1* KO vs WT) revealed a total of 1,712 differentially expressed genes (DEG) and 158 differentially expressed proteins (DEG) (FDR < 0.05; **Figure 1C, D**). Although the number of DEG detected in the proteome was 10 times lower than in the transcriptome, a similar gene expression profile was observed in both data sets **(Figure 1C, D).** Hence, despite the different resolution of these techniques, they are consistent with each other **(Figure 1B)**.

### RNA-seq profiling of *ALMS1*-deficient BJ-5ta cells reveals different pathways related to extracellular matrix regulation

In the transcriptome, of 1,712 DEG, 1,119 (65%) were under-expressed and 593 (35%) were over-expressed in the *ALMS1* KO cell line compared to controls **(Figure 2A; Table S1)**.

**Figure 2.**
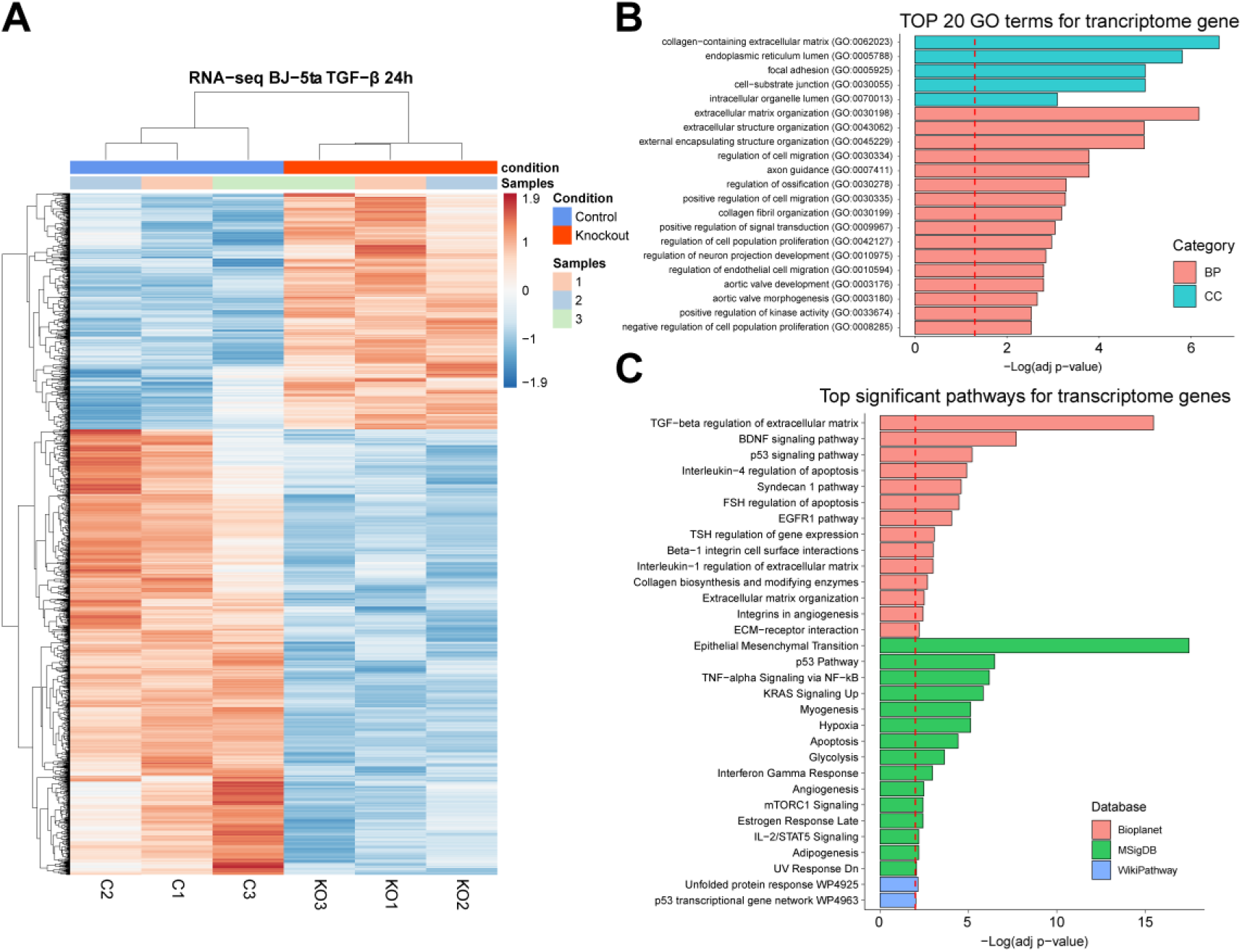
RNA-seq expression profile and enrichment analysis in the ALMS1 knockout cell line (hTERT-BJ-5ta). **(A)** Heatmap of the 1,712 differentially expressed genes with FDR < 0.05 in the hTERT-BJ-5ta cell line after 24 hours with TGF-β pathway stimulation **(B)** The 20 most statistically significant GO terms detected after over-representation analysis (ORA) in the transcriptome. BP: biological process; CC: cellular Component **(C)** Signalling pathways significantly enriched (ORA, FDR < 0.01) for trancriptomic data in Bioplanet, KEGG, MSigDB and Wikipathway databases.

To determine the processes affected by inhibition of *ALMS1* gene expression in the transcriptome, we performed an over-representation analysis (ORA) **(Table S3)**. DEGs were not filtered by fold change (FC) due to the additive effect of small FCs along a signalling pathway **(Methods)**.

We found changes in the gene expression profile of *ALMS1* KO cell line in pathways related to extracellular matrix (ECM) regulation through TGF-β, the Epithelial Mesenchyme Transition (EMT) and various pathways that cross-signal with the TGF-β pathway (BDNF, p53, TNF-α, EGFR1, KRAS and mTORC1) **(Figure 2B, C)**.

Next, we constructed a network using EnrichmentMap (Merico et al., 2010) to reduce the redundancy of pathways and processes between the different databases used in the previous analysis. For this, only enrichR terms with an FDR < 0.05 were used **(Methods)**. In this network, we found two main clusters, highly connected to each other, incorporating pathways from different databases. The most representative keywords in these clusters pointed towards collagen and ECM regulation and β-oxidation of fatty acids **(Figure 3)**.

**Figure 3.**
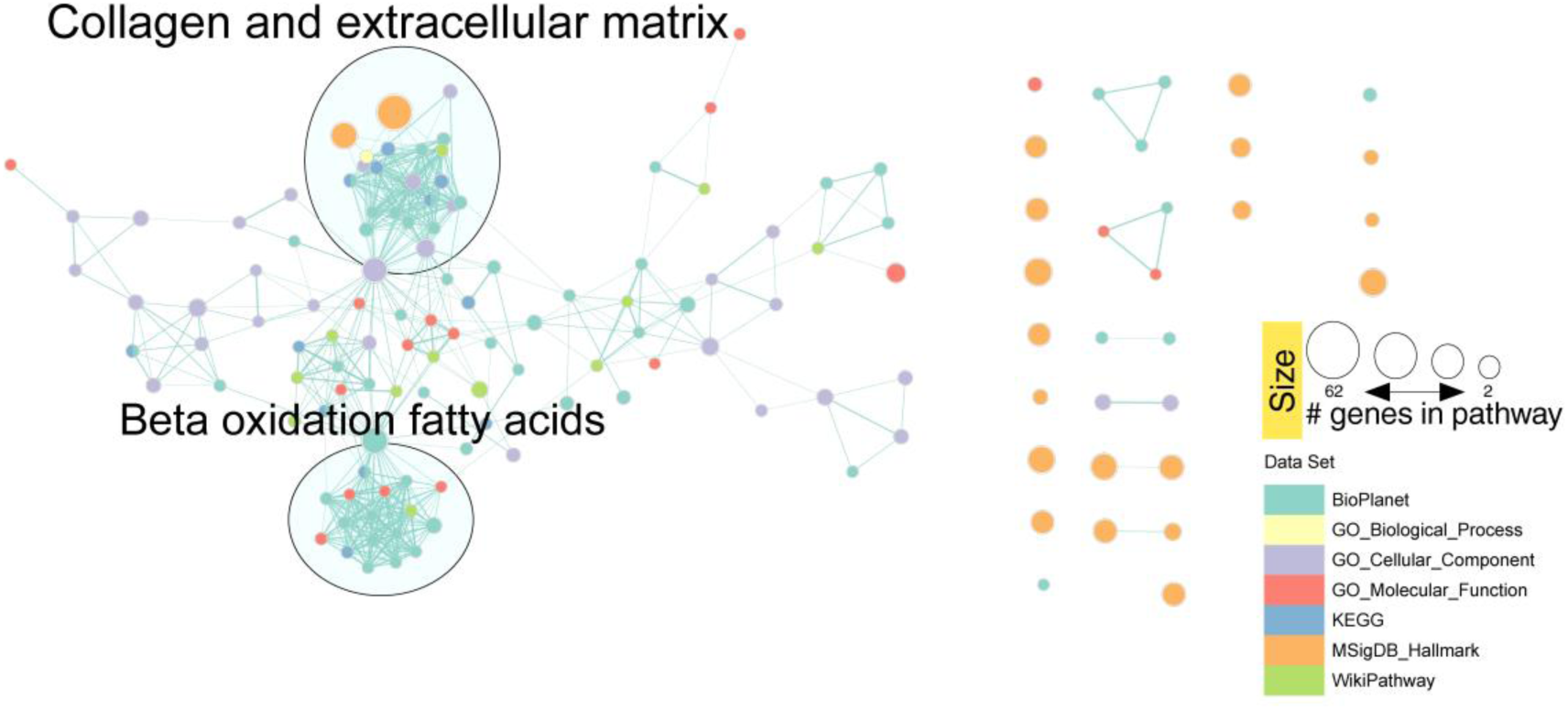
Network clustering of transcriptomic GO terms and enriched routes in the different databases after applying the Markov clustering algorithm (MCL).

### Proteomic profiling of *ALMS1*-deficient BJ-5ta cells suggests alterations in intracellular organelles lumen and lipid metabolism

To study whether the observed differences in gene expression profile are replicated at the protein level, we performed the same analysis on the proteome **(Table S2)**. Of 158 DEG, 98 (62%) were under-expressed and 60 (38%) were over-expressed in the *ALMS1* KO cell line **(Figure 4A)**.

**Figure 4.**
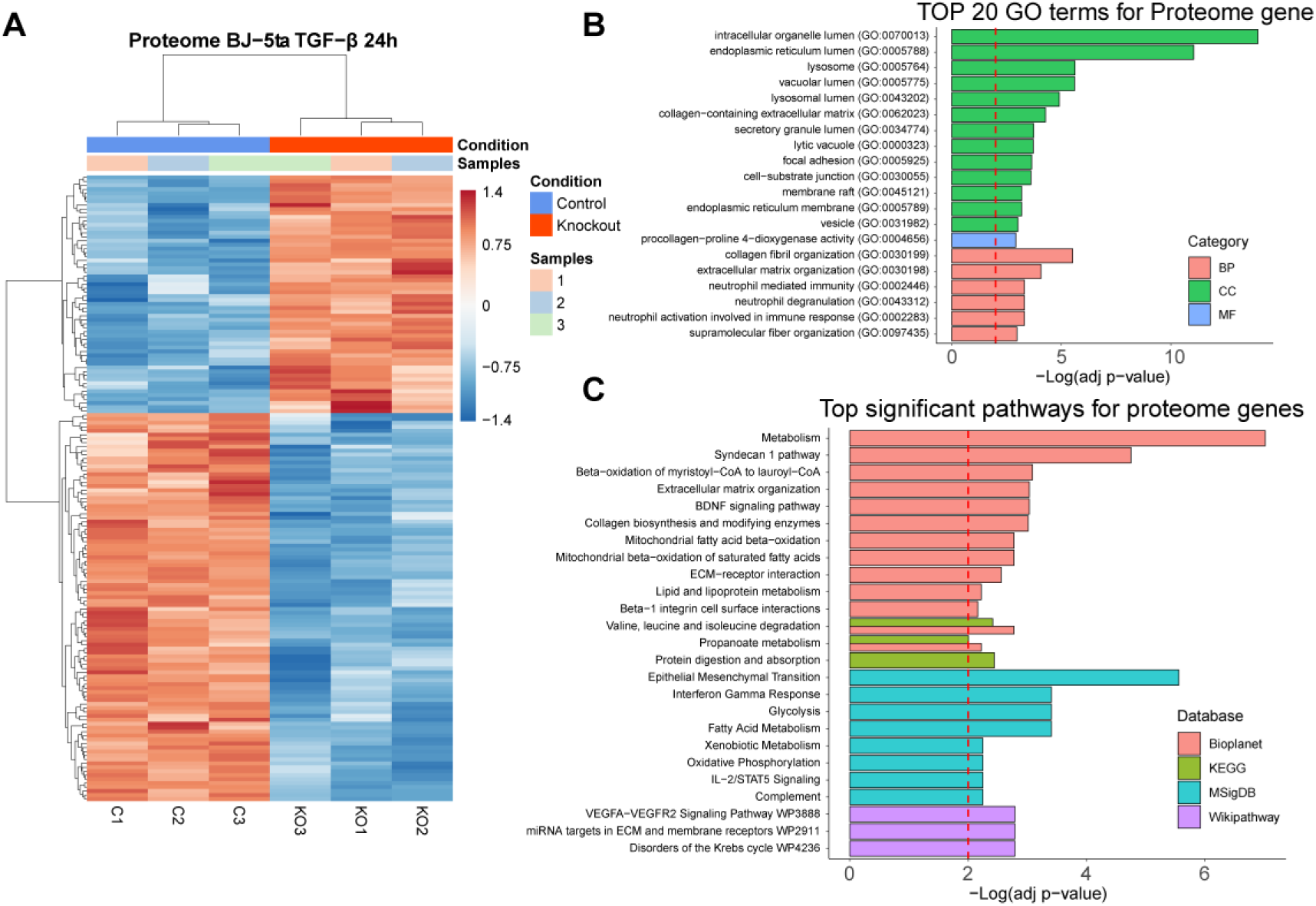
Protein expression profiling and enrichment analysis of ALMS1 knockout cell line (hTERT-BJ-5ta). **(A)** Heatmap of the 158 differentially expressed protein coding genes with FDR < 0.05 in the hTERT-BJ-5ta cell line after 24 hours with TGF-β pathway stimulation **(B)** The 20 most statistically significant GO terms detected after over-representation analysis (ORA) in the proteome. BP: biological process; CC: cellular Component; MF: molecular function **(C)** Signalling pathways significantly enriched (ORA, FDR < 0.01) for proteomic data in Bioplanet, KEGG, MSigDB and Wikipathway databases.

We performed an ORA to establish the proteome-enriched pathways and GOterms **(Table S4)**. The proteome GO terms suggested that depletion of *ALMS1* gene affects the endoplasmic reticulum, lysosomes, focal adhesion and the organisation of collagen and ECM regulation **(Figure 4B)**. These results are consistent with transcriptome findings, including BDNF, Syndecan-1 signalling pathways and EMT **(Figure 4C)**. In addition, other enriched proteomic pathways involved in metabolic processes such as β-oxidation of fatty acids in mitochondria were also detected **(Figure 4C)**.

Regarding the reduction of redundancy between the databases used for the enrichment analysis, we applied the same methodology as in the previous case, obtaining consistent results with the transcriptome. We found two main clusters, associated with collagen and ECM regulation and β-oxidation of fatty acids **(Figure 5)**.

**Figure 5.**
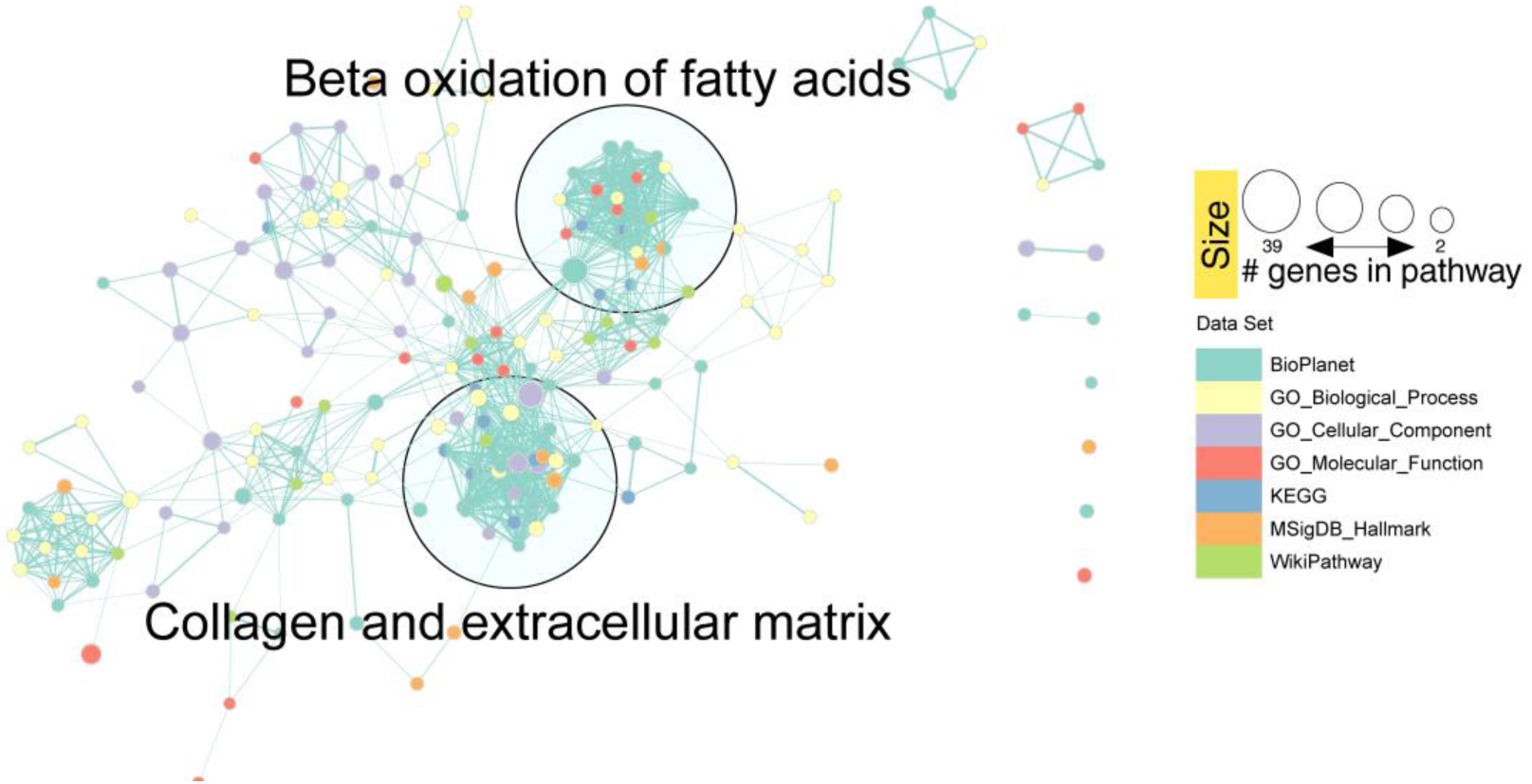
Network clustering of proteomic GO terms and enriched routes in the different databases after applying the Markov clustering algorithm (MCL).

### The multi-omics analysis highlights the association of ALMS1 with the endoplasmic reticulum, Syndecan-1 pathway and EMT

To investigate the relationship between the transcriptome and proteome, we analysed the overlap of DEG between them (1,712 genes and 158 proteins). This resulted in 75 DEG in the *ALMS1* KO cell line **(Figure 6A; Table S5)** with a Pearson correlation of 0.75 (p-value 1.3×10^-14^) between the FCs of the transcriptome and proteome **(Figure 6B)**. Importantly, the likelihood of finding by chance 75 or more common DEG between the transcriptome and proteome is practically zero (Fisher’s exact test). Thus, these genes are a valid signature of ALMS1 depletion **(Figure 6C)**.

**Figure 6.**
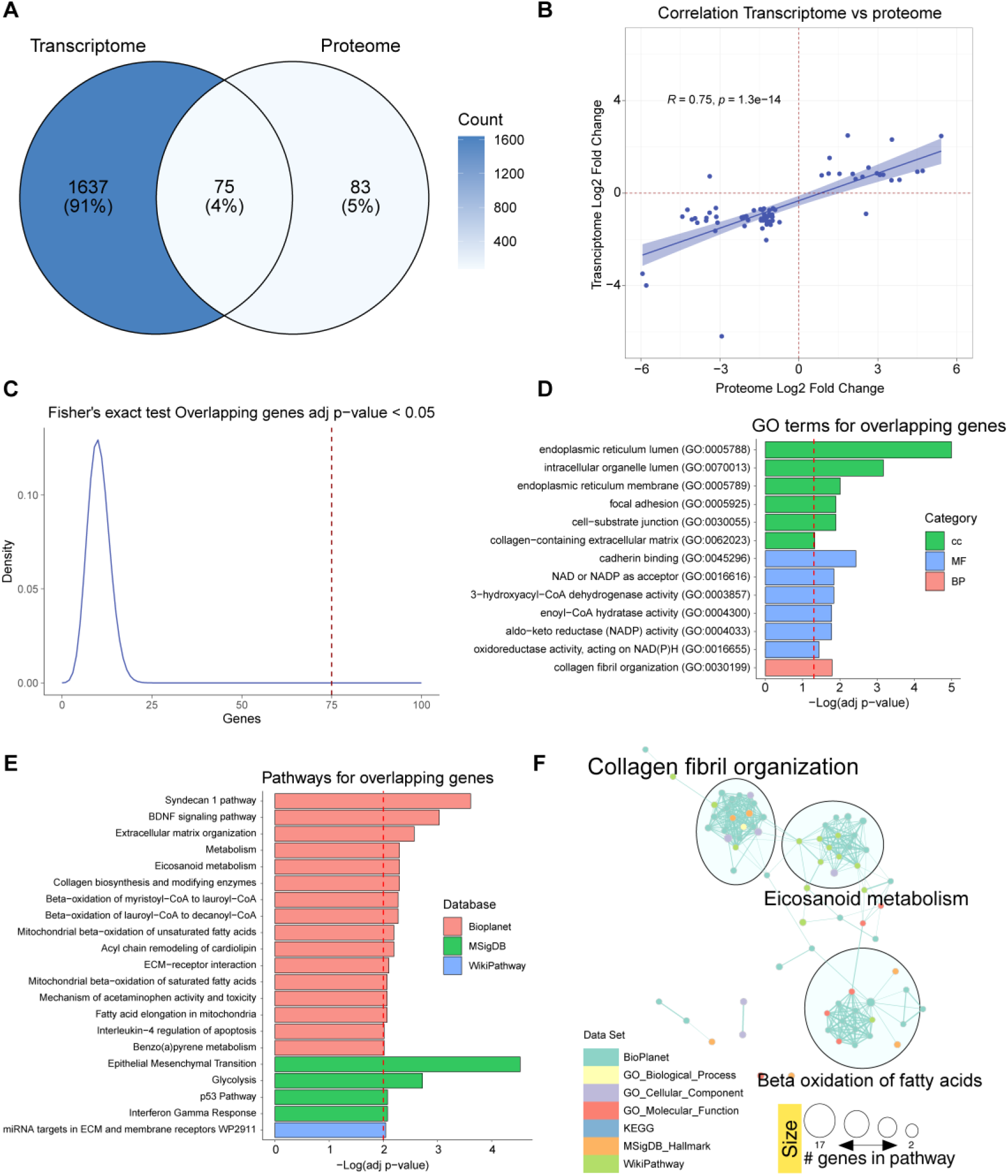
Analysis of differentially expressed genes overlapping between transcriptome and proteome. **(A)** Venn diagram with differentially expressed genes in the transcriptome and proteome and matching terms in both datasets. **(B)** Correlation plot between Log2 FCs in the transcriptome and the proteome of the matched terms. **(C)** Fisher’s exact test to evaluate the probability of obtaining 75 or more matched genes. **(D)** Significantly over-represented (ORA, FDR < 0.05) GO terms that are associated with the subset of matched data. **(E)** Significantly over-represented pathways (ORA, FDR < 0.01) in the list of overlapping genes **(F)**. Network clustering of GO terms and pathways in the different bases of the overlapping gene data after applying the Markov clustering algorithm (MCL).

Further, their enrichment analysis highlights the role of ALMS1 in the endoplasmic reticulum, BDNF or Syndecan 1 pathways, EMT and lipid-associated signalling pathways such as fatty acid metabolism **(Figure 6D, E; Table S6)**. This time, the enriched pathway reduction analysis identified three clusters, one cluster agglutinated the alterations of the ECM, another the alterations of the β-oxidation of fatty acids and the third cluster highlighted the metabolism of eicosanoids, which also affects lipid metabolism **(Figure 6F)**.

### ALMS1 depletion affects the signalling cross-talk between TGF-β and PI3K/AKT

ALMS1 depletion does not appear to affect the activation of the canonical TGF-β pathway in all cell types (Bea-Mascato et al., 2022) so the observed alterations could be due to aberrant signalling of non-canonical pathways such as PI3K/AKT. The PI3K/AKT pathway regulates many of the altered processes found in our analyses. This pathway is involved in control of cell migration, proliferation and adhesion (Xu et al., 2015; Yu and Cui, 2016) as well as in cell metabolism (Yu and Cui, 2016; Hoxhaj and Manning, 2019). Given the present results, we decided to study the cross-signalling between the TGF-β and PI3K/AKT pathway in our *ALMS1*-deficient fibroblast line.

For this purpose, AKT protein phosphorylation levels were measured in the hTERT-BJ-5ta *ALMS1* KO cell line by Western blot at four different stimulation times with TGF-β1 ligand (0, 10, 30 and 90 min). Indeed, deficiency of *ALMS1* generates an over-activation of the AKT pathway **(Figure 7A, B)**. This over-activation was present at basal level (time 0), but was exacerbated by TGF-β stimulation, with the differentials between ALMS1KO and WT being most notable in AKT after 30min of TGF-β1 stimulation **(Figure 7A, B).**

**Figure 7.**
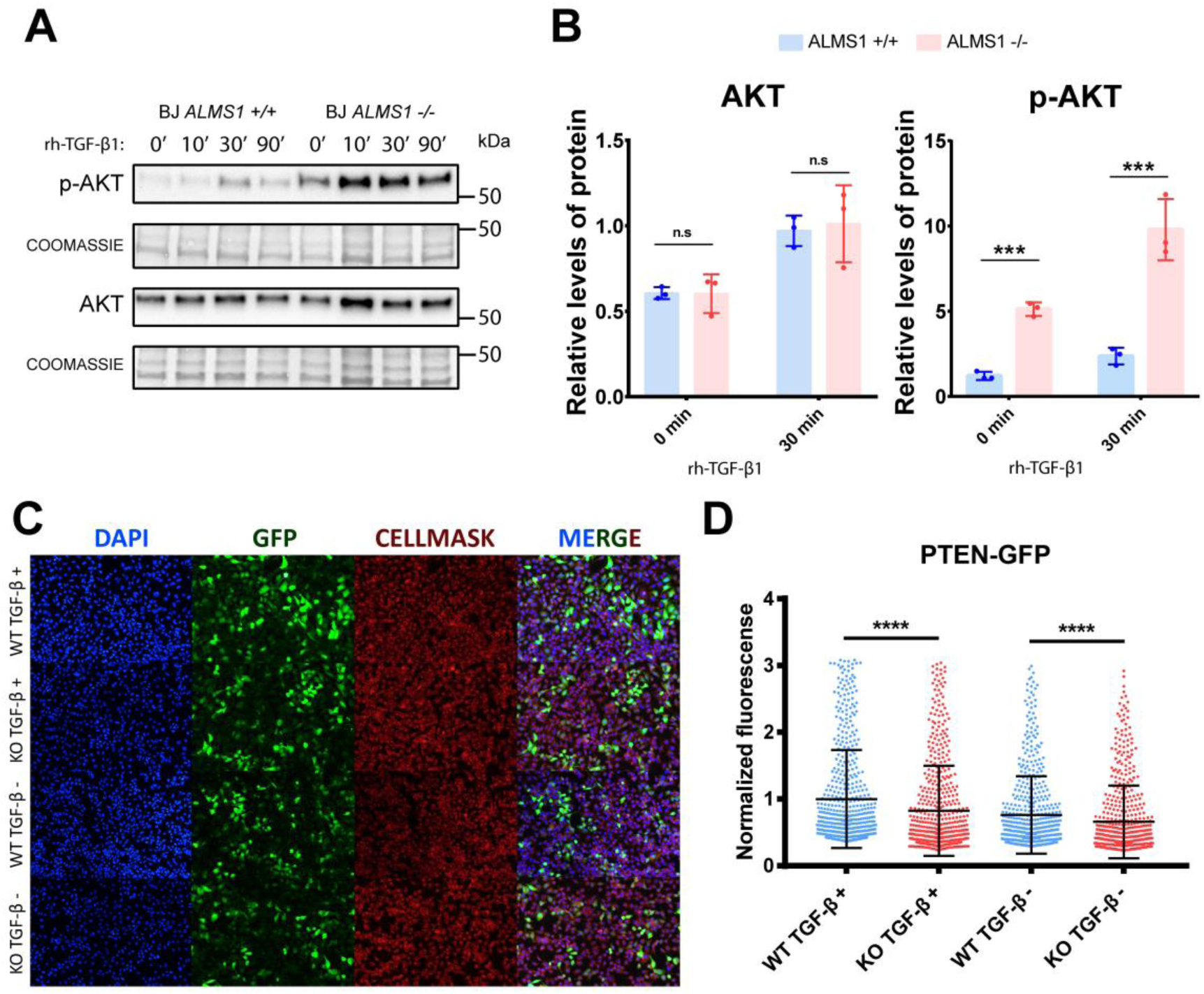
Study of cross-signalling between TGF-β and AKT pathways in ALMS1KO cell models. **(A)** Representative Western blot of AKT overactivation after TGF-β pathway stimulation in BJ-5ta. **(B)** Quantification of AKT protein expression and p-AKT activation levels normalised against Coomassie staining in BJ-5ta. **(C)** Representative images of fluorescence labelling of HeLa cells after transfection with GFP-tagged PTEN. **(D)** Normalised fluorescence of the 550 most fluorescent HeLa cells in each preparation.

We thus hypothesised that over-activation of the AKT pathway results from the accumulation of membrane phospholipids such as PIP3, which is the substrate for the activation of PI3K and subsequently AKT. The PTEN protein is responsible for dephosphorylating PIP3 to PIP2 which inhibits the AKT pathway. For this reason, the PTEN inhibition could be a cause of the AKT over-activation. To test this hypothesis, we transfected a KO model for the *ALMS1* gene in the HeLa cell line with a plasmid to express PTEN protein fused to GFP. We observed a significant reduction of GFP fluorescence in the KO cells compared with controls in both untreated and TGF-β ligand-treated cells (p-value < 0.0001). This suggests that ALMS1 depletion affects PTEN expression. Moreover, this inhibition seems to be slightly accentuated (p-value < 0.01) after TGF-β stimulation **(Figure 7C, D)**.

## DISCUSSION

While the role of the centrosome in the control of the mitotic cycle has been very well studied, little is known about its structure and functions in other contexts (Doxsey et al., 2005; Bettencourt-Dias and Glover, 2007; Conduit et al., 2015). The study of different mutated centrosomal proteins and their impact on processes of the primary cilium, such as ciliogenesis, endocytosis or the signalling of various pathways such as WNT, SHh or TGF-β, has however been enlightening (Kobayashi and Dynlacht, 2011; Collin et al., 2012; Hehnly et al., 2012; Leitch et al., 2014; Mönnich et al., 2018; Gonçalves et al., 2021). Despite all this, many of the regulatory aspects of how centrosomal proteins coordinate these processes remain unknown.

The TGF-β signalling is one of the main pathways regulated through the primary cilium (Clement et al., 2013). It is involved in normal development in mammals and influences other pathways by cross-signalling such as p53, MAPKs or PI3K/AKT (Rahimi and Leof, 2007; Patel et al., 2019; Finnson et al., 2020). For this reason, it is timely to determine the role of different ciliary and basal body genes, such as *ALMS1,* in their regulation.

Thus, we further explored the previously described association between ALMS1 deficiency and alterations in TGF-β-mediated signal transduction (Álvarez-Satta et al., 2021; Bea-Mascato et al., 2022). We performed gene expression profiling by RNA-seq and LFQ-proteomics in an ALMS1KO hTERT-BJ-5ta cell line stimulated with TGF-β1. This cellular model was previously generated and characterised in our lab (Bea-Mascato et al., 2022). We found that the lack of ALMS1 generates an inhibited gene expression profile after TGF-β1 stimulation, both in the transcriptome and proteome **(Figure 2A; 4A)**. This gene expression profile is consistent with our previously described proteomic results in HeLa and with the basal gene expression profile described in patient fibroblasts (Zulato et al., 2011; Bea-Mascato et al., 2022). This means that the lack of ALMS1 is likely to have an inhibitory impact on most of the processes controlled downstream of this gene, such as mitosis, cell migration or receptor recycling. (Zulato et al., 2011; Leitch et al., 2014; Shenje et al., 2014; Bea-Mascato et al., 2022).

The ORA of transcriptome revealed alterations in the TGF-β signalling pathway and other tangentially regulated pathways such as mTOR, EGFR or p53 and processes such as ECM regulation or EMT **(Figure 2B, C)**. The existence of this cross-signalling is known, however, this is the first time that the impact of ALMS1 depletion on it has been shown, which could open up new avenues of treatment for ALMS patients. (Kato et al., 2009; Hamidi et al., 2017; Patel et al., 2019). On the other hand, It was previously established, that ALMS1 depletion led to aberrant expression of several TGF-β1-induced EMT markers in HeLa and hTERT-BJ-5ta cells (Bea-Mascato et al., 2022). However, these results further establish the connection between ALMS1 depletion and EMT alterations. EMT inhibition supports the poor migration capacity of cells lacking ALMS1 which may be due to cytoskeleton collapse, another event described in cells lacking ALMS1 (Zulato et al., 2011; Collin et al., 2012; Bea-Mascato et al., 2022).

Downstream activation of the TGF-β pathway can follow *via* the canonical pathway, dependent on SMADs (SMAD2 and SMAD3), or non-canonical, which encompasses a plethora of pathways such as MAPKs, ERKs, p53, or PI3K/AKT (Finnson et al., 2020). For this reason, alterations in cross-signalling pathways such as mTORC1, EGFR1 or p53 detected in the transcriptome show that alterations in TGF-β are mainly related to the non-canonical part of this pathway **(Figure 2C)** (Patel et al., 2019; Finnson et al., 2020).

Our proteomic analysis revealed proteins involved in lipid metabolism and cytoplasmic organelles such as the endoplasmic reticulum, mitochondria, or lysosomes **(Figure 4B, C)**. We have previously described that ALMS1 depletion does impact cell viability due to changes in mitochondrial reduction capacity (Bea-Mascato et al., 2022). However, it appears to affect mitochondrial β-oxidation of fatty acids **(Figure 4C)**. This could connect the absence of ALMS1 to oxidative metabolism disorders (Liu and Desai, 2015). In addition, alterations in cytoplasmic organelles, such as the endoplasmic reticulum and lysosomes, could be related to alterations in endocytosis, receptor recycling or autophagy. ALMS1 absence has been linked to alterations in transferrin (TfR), NOTCH and NKCC2 receptor recycling, but its role in TGF-β receptor recycling or autophagy has not yet been established (Collin et al., 2012; Leitch et al., 2014; Jaykumar et al., 2018; Yang et al., 2020).

Indeed, after applying a redundancy reduction analysis of enriched pathways, we found that ALMS1 regulates two main clusters in the transcriptome and proteome; processes related to the regulation of the ECM and processes related to lipid metabolism such as β-oxidation of fatty acids **(Figure 3; 5)**. It has already been described that deletion of ALMS1 leads to overexpression of several ECM components in fibroblasts from ALMS patients. For this reason, understanding these alterations is key to determining the pathophysiology of fibrosis in ALMS (Zulato et al., 2011). On the other hand, the metabolic alterations observed in ALMS could have their origin in processes such as β-oxidation of fatty acids (Romano et al., 2008; Huang-Doran and Semple, 2010; Favaretto et al., 2014; Geberhiwot et al., 2021).

The list of common differentially expressed genes between the transcriptome and proteome supports the above findings, indicating that ALMS1 depletion affects EMT, endoplasmic reticulum, Syndecan-1 and BDNF signalling pathways and lipid metabolism pathways such as β-oxidation of fatty acids **(Figure 6D, E)** (Kasza et al., 2014). By studying this list of genes by redundancy reduction analysis, we saw the existence of a link between ALMS1 depletion and eicosanoid metabolism in addition to the processes mentioned above **(Figure 6F)**. Alterations related to the ECM are undoubtedly key to the appearance of fibrosis in these patients; however, alterations in the β-oxidation of fatty acids, beyond their link to obesity, may contribute to the appearance of fibrotic tissue due to lipid deposits (Serra et al., 2013; Kang et al., 2014). This suggest that the metabolic disorder in patients with ALMS is involved in the progressive development of fibrosis in various organs such as the kidneys and liver (Bettini et al., 2021).

Regarding the deregulation of the PI3K/AKT, we can relate the over-activation of AKT **(Figure 7A, B)** with the inhibition of the p53 pathway **(Figure 1C)** (Gottlieb et al., 2002; Abraham and O’Neill, 2014). This is supported by the decrease in the expression of the *PTEN* gene, which could explain an over-phosphorylation of the cell membrane and an accumulation of PIP3 leading to the over-activation of AKT **(Figure 7C, D)** (Álvarez-Garcia et al., 2019). To our knowledge, this is the first study to explore how *ALMS1* affects cross-signalling between AKT and the TGF-β pathway. Other studies have been carried out to try to understand the role of *ALMS1* in regulating the insulin mediated AKT pathway. The involvement of AKT has not yet been clearly established in ALMS, while some studies in pre-adipocytes have not detected any differences in AKT activation, other studies in mice have shown that AKT is over-activated in some tissues and it is inhibited in others, when *ALMS1* is depleted (Huang-Doran and Semple, 2010; Favaretto et al., 2014; Geberhiwot et al., 2021). AKT regulation is tissue specific, it has 3 distinct isoforms and each has multiple phosphorylation sites (Liao and Hung, 2010). This could be an explanation for the disparity of the data present in the bibliography.

The over-activation of the AKT pathway could be the link between alterations at the metabolic level and alterations related to the ECM. The inhibition of p53 would be caused by this over-activation through MDM2, affecting the PTEN-p53-AKT-MDM2 loop (Gottlieb et al., 2002; Abraham and O’Neill, 2014). This hypothesis would be supported by the resistance to apoptosis in *ALMS1*-depleted cells (Zulato et al., 2011; Bea-Mascato et al., 2022).

## CONCLUSION

In conclusion, the lack of *ALMS1* gene expression alters the TGF-β pathway in the knockout cell model. It also seems to compromise the signalling of other tangential pathways such as PI3K/AKT or p53. These signalling alterations appear to affect diverse metabolic processes involving the mitochondrial, endoplasmic reticulum and lysosomes and various lipid-mediated signalling pathways. The alterations are mainly clustered into processes affecting the regulation of the ECM and the β-oxidation of fatty acids, which we confirmed in both the transcriptome and proteome. Finally, *ALMS1* depletion led to over-activation of the AKT protein in BJ-5ta cells and inhibition of the PTEN protein in HeLa cells.

## Author Contributions

BB-M, DV designed the study. BB-M performed the experiments, analysed the data, interpreted the results, created the figures, and drafted the first manuscript. EG-C, YS-C and SC helped with RNA-seq analysis and data integration. DV supervised the study and performed funding acquisition. All authors provided comments and corrections to the manuscript and gave their approval for publication.

## Informed Consent Statement

Not Apply.

## Data Availability Statement

Data are available via Gene Expression Omnibus (GEO) with the identifier GSE209844 and ProteomeXchange with the identifier PXD035708.

## Code Availability Statement

The code used in this study can be accessed at the GitHub address “https://github.com/BreisOne/multiomicpipeline”.

## Acknowledgements

We sincerely thank the Proteomics and Genomics services from “Centro de Apoyo Científico-Tecnológico a la Investigación” (CACTI) of University of Vigo and its specialists Paula Álvarez Chaver, Ángel Sebastián Comesaña, Verónica Outeiriño and Manuel Marcos for their guidance and advise. We also thank Mercedes Peleteiro Olmedo from “Centro de Investigacións Biomédicas” (CINBIO) from University of Vigo for the flow cytometry service.

