## Supplementary Figures S1-2 for "Loss of the centrosomal protein ALMS1 alters lipid metabolism and the regulation of extracellular matrix-related processes"

### Supplementary Figures:

**A**

| Sample Name | % Assigned | M Assigned | % Aligned | M Aligned | % Dups | % GC | Length | M Seqs |
|---|---|---|---|---|---|---|---|---|
| C1 | 15.6% | 11.6 | 47.6% | 12.5 | 66.8% | 52% | 122 bp | 26.3 |
| C2 | 20.1% | 14.2 | 51.1% | 15.3 | 70.8% | 53% | 140 bp | 29.8 |
| C3 | 19.0% | 10.8 | 48.9% | 11.6 | 63.6% | 51% | 100 bp | 23.6 |
| KO1 | 16.1% | 19.0 | 46.0% | 20.4 | 62.3% | 54% | 111 bp | 44.4 |
| KO2 | 20.6% | 11.5 | 53.5% | 12.3 | 52.0% | 52% | 95 bp | 23.1 |
| KO3 | 24.5% | 16.1 | 56.8% | 17.2 | 63.1% | 53% | 125 bp | 30.3 |

**B**

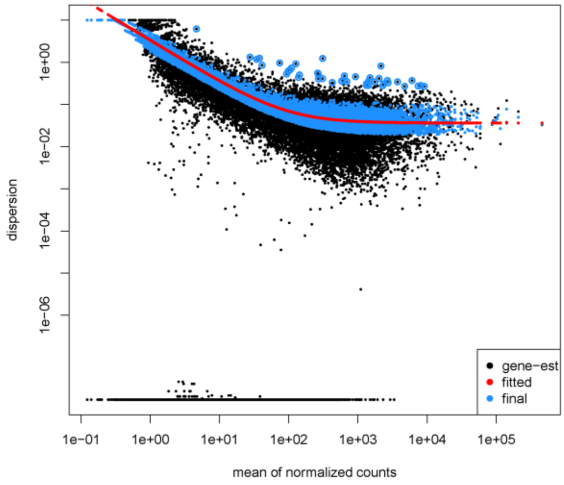


**C**

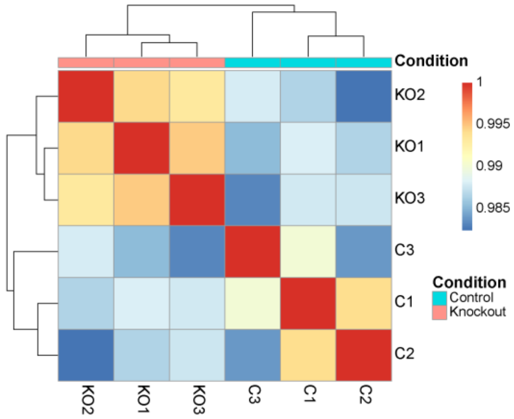


**D**

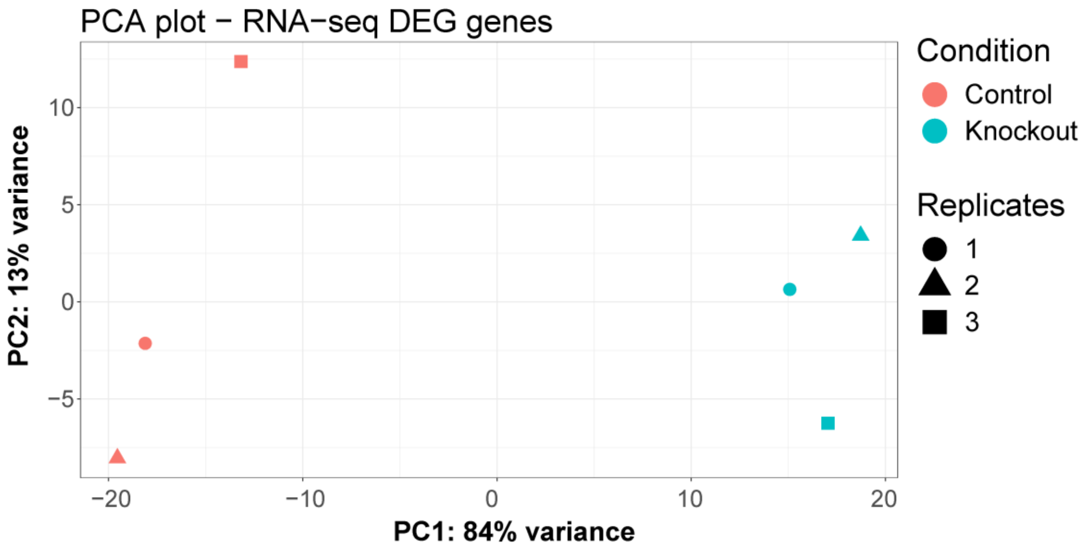

**Supplementary Figure S1.** RNA-seq quality controls. **(A)** Table with the quality controls, showing percentage of assigned, aligned and duplicated reads, GC percentage and average read length. **(B)** Dispersion plot showing the decrease in variance as the normalised mean number of counts in each gene increases. **(C)** Correlation matrix showing similarity between biological replicates of each genotype. **(D)** PCA plot of the RNA-seq samples, where PC1 (genotype) accounts for 84% of the variance between samples.

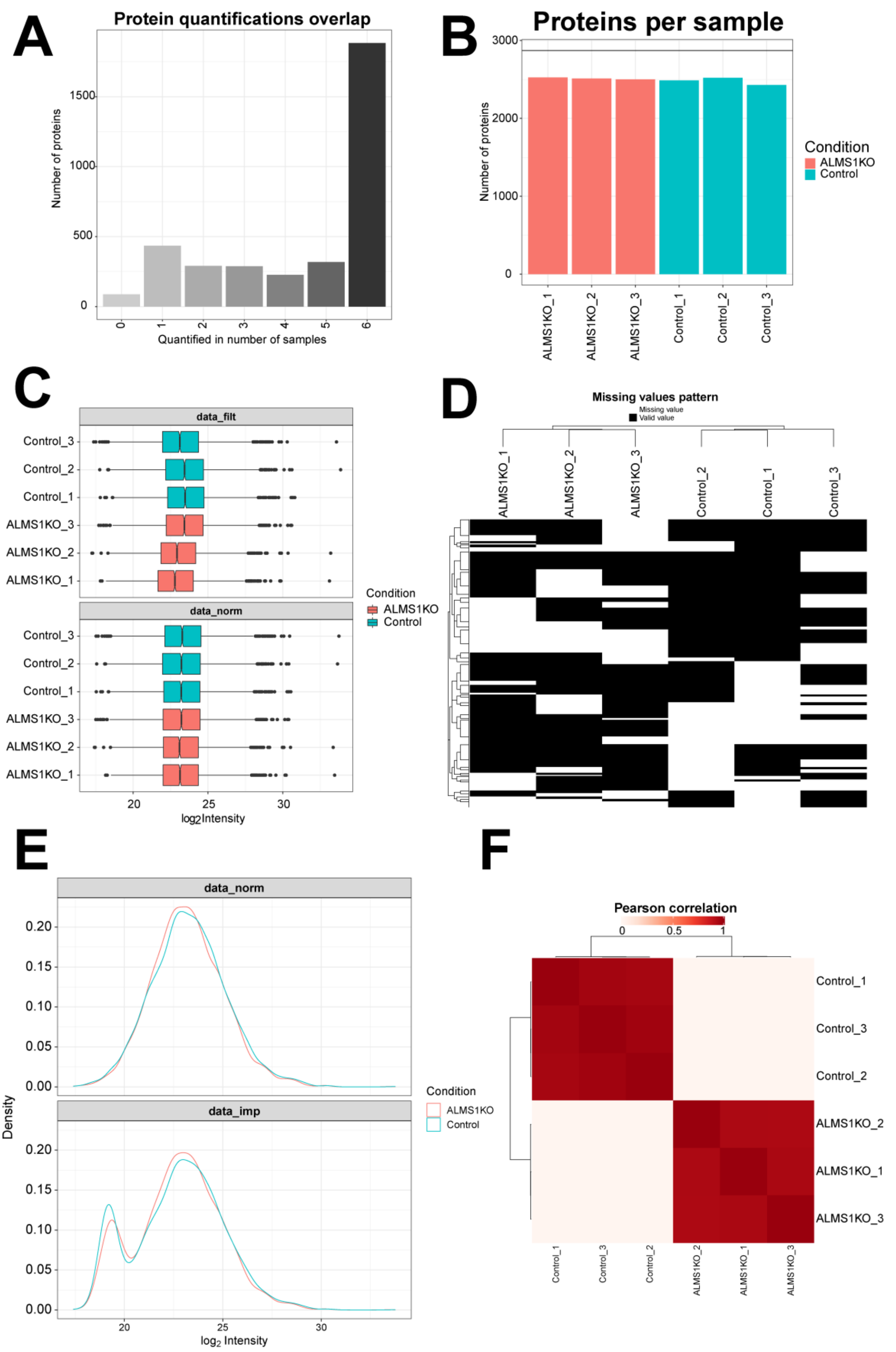


**Supplementary Figure S2.** Proteomics quality controls **(A)** Overlap of proteins quantified in each sample. A total of 1,800 proteins were quantified in all samples. **(B)** Number of proteins identified in each sample. Approximately 2,870 proteins were identified in each sample. **(C)** Box-plots showing the distribution of log2 Intensity of each protein per sample before and after applying VSN normalisation. **(D)** Heatmap showing the pattern of non-random missing values prior to imputation. **(E)** Density-plot of the distribution of the average log2 values of the intensities between WT and KO before and after applying the Minprob imputation with an FDR > 0.01. **(F)** Correlation matrix showing similarity between biological replicates of each genotype.

### Original Western Blots:

### p-AKT:

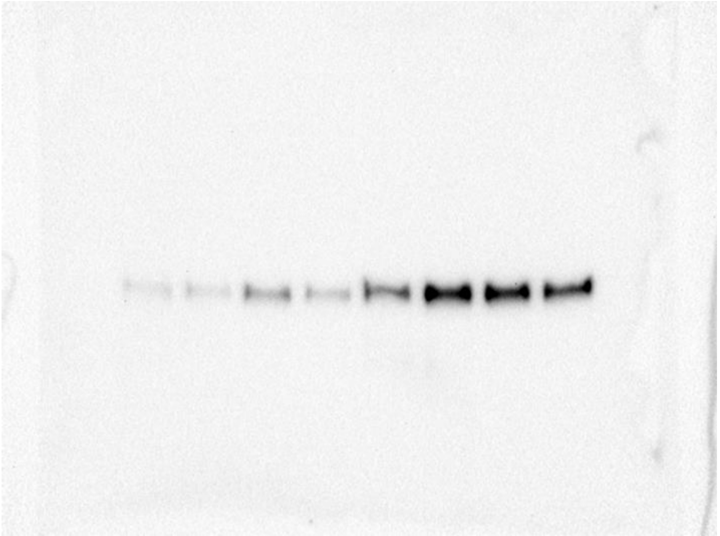

### AKT:

The first well was a bad load of WT time 0.

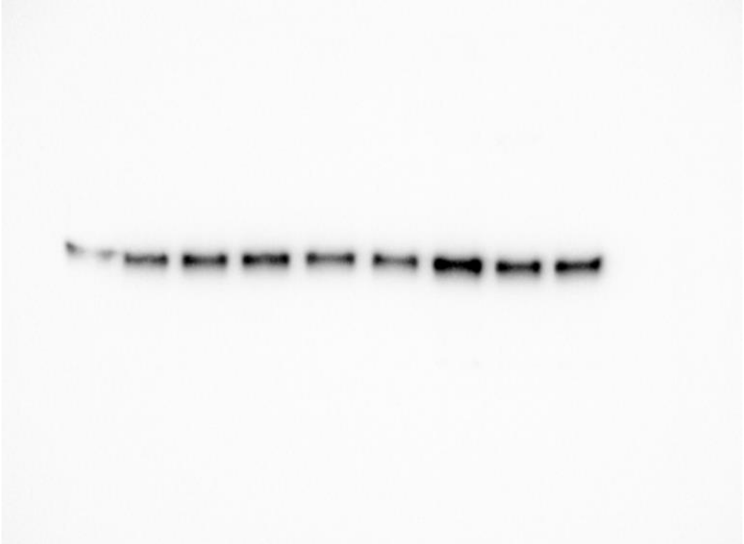
